# PhysiGym: bridging the gap between the Gymnasium reinforcement learning application interface and the PhysiCell agent-based model software

**DOI:** 10.1101/2025.09.18.677030

**Authors:** Alexandre Bertin, Elmar Bucher, Owen Griere, Marcelo Hurtado, Heber Rocha, Randy Heiland, Aneequa Sundus, Paul Macklin, Vincent François-Lavet, Emmanuel Rachelson, Vera Pancaldi

## Abstract

This paper presents PhysiGym, a framework that integrates agent-based biological simulation within standardized reinforcement learning environments. By integrating the agent-based modeling framework PhysiCell with the Gymnasium API, we provide a flexible tool for exploring reinforcement learning strategies to control insilico biological processes. We demonstrate PhysiGym’s potential with a case study where a deep reinforcement learning algorithm guides a tumor microenvironment model toward an anti-tumoral state, ultimately achieving tumour elimination. Our results highlight PhysiGym’s flexibility for AI-driven biological control and optimization of dynamic treatment regimes.

## 1 Introduction

Biological systems are complex, dynamic, and often difficult to study in real life. In silico modeling aims to mimic these systems, providing a controlled environment for exploring specific aspects of biological processes and testing hypotheses. In particular, agent-based models (ABMs) provide a powerful framework for simulating biological systems where agents (e.g. cells) interact dynamically based on predefined rules. These models are used in oncology and immunology to study emerging behaviors in complex biological systems, with the potential to propose novel therapeutic strategies [9, 10, 18].

Ordinary differential equations (ODEs) and agent-based models (ABMs) are widely used to simulate the tumor microenvironment (TME). ODEs provide compact system-level descriptions, while ABMs capture spatial and cellular heterogeneity at higher computational cost. For dynamic treatment regimes (DTRs), however, simulation alone is insufficient: designing adaptive, sequential interventions requires a control framework. Reinforcement learning (RL), where an RL agent learns policies through interaction with an environment, offers such a framework [2]. By modeling treatment regimes as a sequence of decisions, RL enables adaptive control of biological simulations [4]. For efficient experimentation and policy learning, RL further requires a standardized interface. To address this, we introduce PhysiGym, an interface that connects PhysiCell [6], an ABM framework for multicellular simulations of tissues or cell cultures (tumor microenvironment, other disease tissues, or bacteria colonies), with Gymnasium [20], a widely adopted RL application programming interface (API). Physi-Cell enables flexible multiscale modeling of biological systems, coupled with a sophisticated underlying physics simulator (the BioFVM diffusion transport solver [5]), while Gymnasium provides a structured framework for training and evaluating RL policies.

The first implementation of RL on an ABM framework based on BioFVM was proposed by Zade et al. [23], which applied Q-learning [22] to optimize Temozolomide treatment regimes for patients with glioblastoma multiforme, a highly specific problem. More recently, Aif et al. [1] used PhysiCell as a complex (high-fidelity) simulation environment for tumor growth, combined with a simple (low-fidelity) RL training environment of coupled ordinary differential equations (ODE) to learn treatment strategies for solid tumors.

PhysiGym bridges PhysiCell and Gymnasium to provide a reproducible and scalable framework for integrating agent-based simulations of biological systems with RL strategies. In the following, we present PhysiGym’s motivation and design, demonstrating how it directly connects ABMs and RL to control complex biological systems. In particular, we focus on the tumor microenvironment, a biological system made up of cancer cells and surrounding immune and stromal cells that engage in complex interactions and cross-talks, which is proven to be crucial for an effective response to immunotherapy, an important therapeutic approach in an increasing number of types of cancer.

## 2 Methods

### 2.1 PhysiGym design and implementation

PhysiCell is an ABM framework written in C++ and implemented to model multicellular systems based on classical mechanics [6]. Cells are agents whose type determines the specific rule set that governs their behavior and interactions with the other agents and the environment. Substrates, like oxygen or cytokines, can be modeled with the integrated BioFVM diffusive transport solver [5]. In addition, intracellular models can be integrated into cell agents [11, 15, 19] to model specific behaviors based on each cell’s environment and gene regulatory networks. The extracellular matrix that surrounds cells in tissues can also be modeled as a mesh of fibers for a more realistic representation of the cells’ physical environment [13]. Together, these components enable the construction of ABMs that are spatially explicit in two or three dimensions, off-lattice, center-based, and multiscale in both space and time [12].

RL is used to control discrete-time dynamical systems, which are typically modeled as Markov Decision Processes (MDPs) [3, 16]. An MDP consists of four key elements:

1. *S* the state space, where *s* ∈ *S* represents a state of the environment (e.g., a vector or an image).
2. *A* the action space, where *a* ∈ *A* represents an action applied to the environment (e.g. drug dose).
3. *T* the transition model, which defines how the environment evolves. The transition model can be either deterministic or stochastic. Formally, the next state noted *s*^*′*^ is given by *s*^*′*^ ∼ *T* (*s, a*) where *s* represents the current state and *a* the action taken.
4. *R* the reward function, which evaluates whether an action was beneficial. Formally, *r* denotes the reward obtained by taking action *a* in state *s*, i.e., *r* = *R*(*s, a*).

The objective is to find the policy that maximizes the expected discounted cumulative reward:

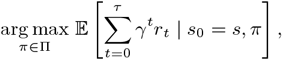

with *s*_0_ the initial state, *π* the policy, *γ* the discount factor, and *τ* the terminal timestep. In the RL community, the Gymnasium RL API is widely used as a standard interface for MDPs, promoting code reuse and modularity in RL algorithms and models.

PhysiGym bridges the gap between the PhysiCell ABM framework and the Gymnasium RL API by extending the Python interpreter [21, 17] with PhysiCell. As a result, the ABM model is implemented in the same way as standard PhysiCell models, using PhysiCell Studio [8] and C++.

Any model implemented in PhysiCell can be converted into a PhysiGym model. The interface between PhysiCell and Gymnasium is based on the PhysiCell “user parameter”, “custom data variable”, and “custom data vectors” data constructs. Information, such as how much drugs to apply to the domain, can only be transferred over these variable types. The variables (for example, drug) be specified in PhysiCell Studio under the `User Params` and `Cell Types / Custom Data` tab. As a minimum requirement, the user must define the parameter *dt_gym*, which specifies the time interval at which interactions with the environment occur, and the parameter *time*, a time tracker necessary to determine when to update the environment. In addition, additional parameters can be defined to control the simulation, such as drug administration, which can serve as actions. Similarly, other parameters can be designated to record specific quantities of interest, which can then be used to construct the observations.

According to the transition function, for each episode (a complete sequence of states, actions, and rewards from the start to the end of a trial), values from these data constructs can be modified without having to recompile the source code. There are three additional functions to define the state space (read information from the simulation): `get_cell` for cell position, `get_microenv` for substrate concentration, and `get_graph` to retrieve cell neighborhood information.

To extend the Python interpreter with PhysiCell, the main simulation loop, written in C++, was split into three functions: `physicell_start`, `physicell_step`, and `physicell_stop`, all of which can be called from Python. The `physicell_start` function initializes the PhysiCell model, while `physicell_stop` terminates the simulation. The `physicell _step` function is responsible for advancing the environment by one time step based on the action taken by the reinforcement learning agent. For most models, only `physicell_step` needs to be modified, e.g. to apply drugs to the ABM. Several additional C++ functions are exposed in Python to facilitate data retrieval. These include functions to obtain custom variables, vectors, `physicell_get_cell`, which provides a matrix containing information about each cell, such as its type, coordinates, and death status, and `physicell_get_microenv`. All exposed functions are implemented in the C++ file physicellmodule.cpp, for more details, consult the reference manual https://github.com/Dante-Berth/PhysiGym/blob/main/man/ REFERENCE.md.

By extending the necessary functions to obtain the desired control over the ABM and modifying the `physicell_step` function, users can compile the C++ code and integrate it with Python, particularly with the Gymnasium API. This allows users to define `physicell_model.py`, which implements a Gymnasium-compatible environment. Specifically, `physicell_model.py` defines a child class of Gymnasium’s environment class, inheriting from `CorePhysiCellEnv` which is specified in `physicell _core.py. CorePhysiCellEnv` is the main class that takes charge of the use of `physicell_start`, `physicell_stop` and `physicell_step` which are model independent. This class handles interactions with the Gymnasium framework. Users should modify physicell model.py to specify the observation space, action spaces, and the reward, as well as any additional data they wish to store.

For an in-depth understanding, tutorials are provided and can be found at the PhysiGym GitHub home page. Besides, pytest unit tests are included in the code base, and we use GitHub Actions for continuous integration. PhysiGym was extensively tested for compiling and running on Linux, Windows Subsystem for Linux, and macOS. Installation and troubleshooting how-tos, a modeling tutorial, an RL tutorial, and a reference manual can be found at [https://github.com/Dante-Berth/PhysiGym/tree/main/man].

### 2.2 Implementation of a tumor microenvironment toy model 1

We developed a simplified tumor microenvironment (TME) model 1 to illustrate the potential uses of PhysiGym. The model includes four cell types, four diffusible substrates, and a set of simple agent-based rules. At the initial state, the total number of tumor cells is 512. And 128 cell 1 surrounds the tumor. Moreover the total number of cell 1 and cell 2 is fixed at 128.

#### 1. Cell types

- **Tumor cells**: proliferate (5 *×* 10^−5^ min^−1^), apoptosis (1 *×* 10^−6^ min^−1^), immotile, uptake anti- and protumoral factors. Necrosis, motility, and baseline secretion disabled.
- **Cell _1**: non-proliferative, mobile, uptakes pro-tumoral factor (10*/*min). Conditional rules allow secretion of anti-tumoral factor and transformation into cell _2.
- **Cell _2**: non-proliferative, mobile, uptakes drug_1 (10*/*min). Conditional rules allow transformation into cell 1 in presence of drug_1.

#### 2. Substrates

1. **Debris**: produced by dead tumor cells, diffusion 1 *µm*^2^*/*min, no decay. Acts as a signal for debris-related rules.
2. **Drug 1**: decay 0.01*/*min, applied at boundaries, regulates `cell_2` →`cell_1` transformation.
3. **Anti-tumoral factor**: diffusion 1000 *µm*^2^*/*min, decay 0.05*/*min, secreted conditionally by `cell _1`, enhances tumor apoptosis.
4. **Pro-tumoral factor**: diffusion 1000 *µm*^2^*/*min, decay 0.1*/*min, can be suppressed by `cell_1` and enhances tumor survival.

#### 3. Cell Hypothesis Rules

In `tumor` cells: pressure decreases cycle entry from 5e-05 towards 0 with a Hill response, with half-max 5 and Hill power 3. pressure increases necrosis from 0 towards 0.0028 with a Hill response, with half-max 10 and Hill power 3. anti-inflammatory factor decreases apoptosis from 1e-06 towards 0 with a Hill response, with half-max 5 and Hill power 10. pro-inflammatory factor increases apoptosis from 1e-06 towards 0.05 with a Hill response, with half-max 1 and Hill power 2. dead increases debris secretion from 0 towards 0.25 with a Hill response, with half-max 0.2 and Hill power 5. Rule applies to dead cells.

In `cell_1` cells: pressure increases transform to `cell_2` from 0 towards 1 with a Hill response, with half-max 11 and Hill power 4. anti-inflammatory factor decreases pro-inflammatory factor secretion from 10 towards 0 with a Hill response, with half-max 10 and Hill power 4.

In `cell_2 cells: drug_1` increases transform to `cell _1` from 0 towards 1 with a Hill response, with half-max 5 and Hill power 4.

This toy model highlights how the RL framework can control tumor progression by modulating the whole microenvironment, rather than solely targeting cancer cell killing.

### 2.3 Implementation of an RL approach to control the TME simulation

Our overall objective is to achieve complete tumor eradication while minimizing drug usage. The action is *d*_*t*_ ∈ [0, 1] what refers to **drug 1** applied to the system. The action space is defined as:

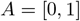

where 0 represents the minimum drug amount, and 1 represents the maximum drug amount added to the tumor microenvironment. The reward function is defined as:

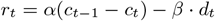

and it consists of two main components: the first term, *α*(*c*_*t*−1_ − *c*_*t*_), promotes a reduction in tumor size. In our model, *α* = 0.0156 is a normalization coefficient that scales the tumor reduction term to be of similar magnitude to the second component, *β* · *d*_*t*_, which penalizes drug usage. The parameter *β* = 0.2 controls the strength of this drug penalty.

Increasing *α* or decreasing *β* leads the reinforcement learning algorithm to prioritize tumor reduction, potentially at the expense of higher drug administration. Conversely, decreasing *α* or increasing *β* encourages the agent to minimize drug usage, even if it means tolerating more tumor growth. In other words, tuning *α* and *β* influences whether the agent prioritizes killing tumor cells or avoiding drug toxicity.

We propose two different observation spaces, depending on whether we consider the spatial aspect of the simulation or not.

The first observation space, called *scalars*, is based on computing the cell count for each cell type and the maximum value for each substrate. We denote by cell_*i*_(*t*) the number of cells of type *i*, with *i* ∈ {1, 2}. Let *c*_*t*_ represent the number of cancer cells at time *t*, and let *c*_init_ denote the initial number of tumor cells (i.e., at *t* = 0). We normalize each quantity by the relative variation of the number of cells with respect to the initial number of cancer cells (arbitrarily set to 512). This is formalized by the function:

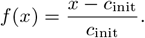

Thus, the observation state at time *t* is given by:

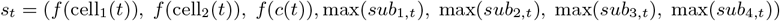

where max(*sub*_*j,t*_) represents the maximum amount of the substrate in the entire environment and *sub*_*j,t*_ denotes substrate *j* at time *t*, with:

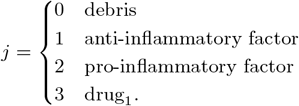

The second observation space, called *multi-channel cells & substrates*, considers the spatial distribution of cells and substrates via an image representation of the simulation state. A set of channels corresponds to a specific cell type, while the other set of channels refers to the substrates. Substrates are chemical products or extracellular proteins produced by cells or produced by the environment or chemical products added to the environment, such as drugs. To reduce memory requirements, we reduce the shape of the original image given by the `PhysiCell_Settings.xml` file by discretizing the continuous environment onto a uniform grid. We also compute 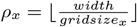 and 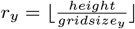. In our environment, *ρ*_*x*_ = *ρ*_*y*_ because *width* = *height* = 512 and *gridsize*_*x*_ = *gridsize*_*y*_ = *gridsize* = 64. The size of the bins is calculated by mapping the continuous coordinates into discrete indices. Specifically:

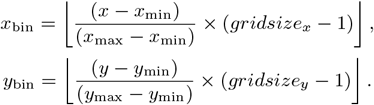

This ensures that the continuous spatial domain is discretized into a grid of size *gridsize × gridsize*. If one or more cells are present in a bin, we increment the count in the channel corresponding to the respective cell type. Formally, for each cell:

- Determine its bin index (*x*_bin_, *y*_bin_).
- Let *c* ∈ 0, 1, 2 be the index associated with its cell type.
- Increment image[*c, y*_bin_, *x*_bin_] by 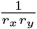.

By dividing by *ρ*_*x*_*ρ*_*y*_, we normalize the count so that the value in each bin represents an area contribution, ensuring that our image values stay in the range [0, 1]. This produces an image tensor of shape (num cell types, *gridsize, gridsize*), Thus, for our three cell types, we can represent the data by images, as shown in figure 2.

**Figure 1.**
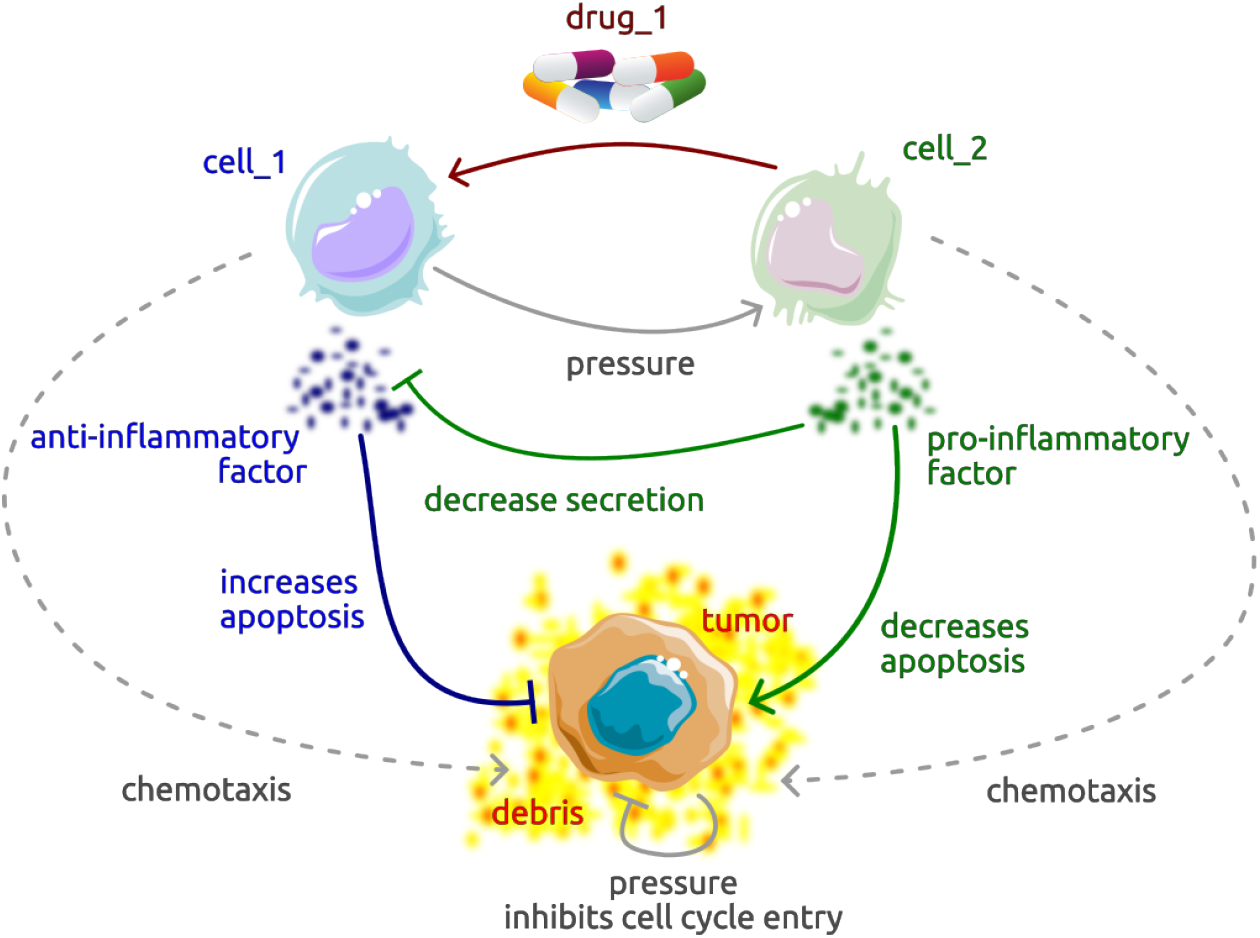
Toy model of the tumor microenvironment, representing interactions between cancer cells and two other distinct cell types with specific behaviors, together with exchanged cytokines and different processes. ⊣ represents an inhibition, while → represents an activation. The color of the arrows corresponds to the color of the cell producing the protein associated with the arrow. For instance, `cell _1` is in blue; thus, the arrow related to increasing apoptosis of the tumor is also in blue, as in `cell_1`.

**Figure 2.**
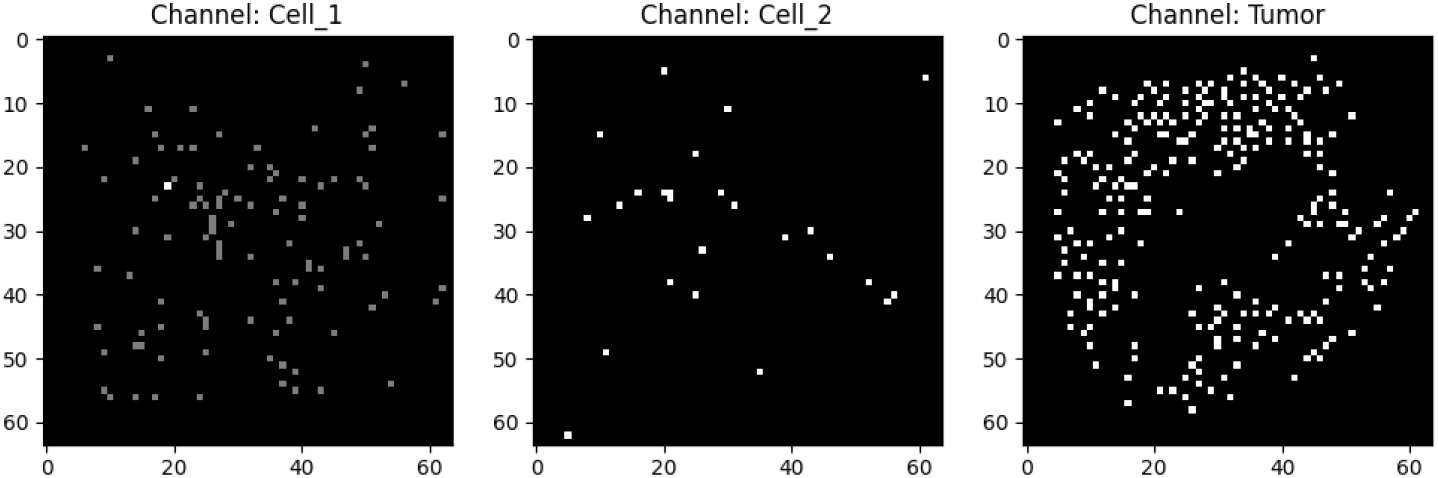
Representation of different channels, each channel represents a cell type.

For substrate channels, the process is similar but simpler, since substrate values are continuous. Using the same (*x*_bin_, *y*_bin_) discretizations, we assign each bin the maximum substrate value observed in that bin. A minmax normalization is then applied across the entire image for each substrate channel, bringing all values into range [0, 1], which can be seen in figure 3.

**Figure 3.**
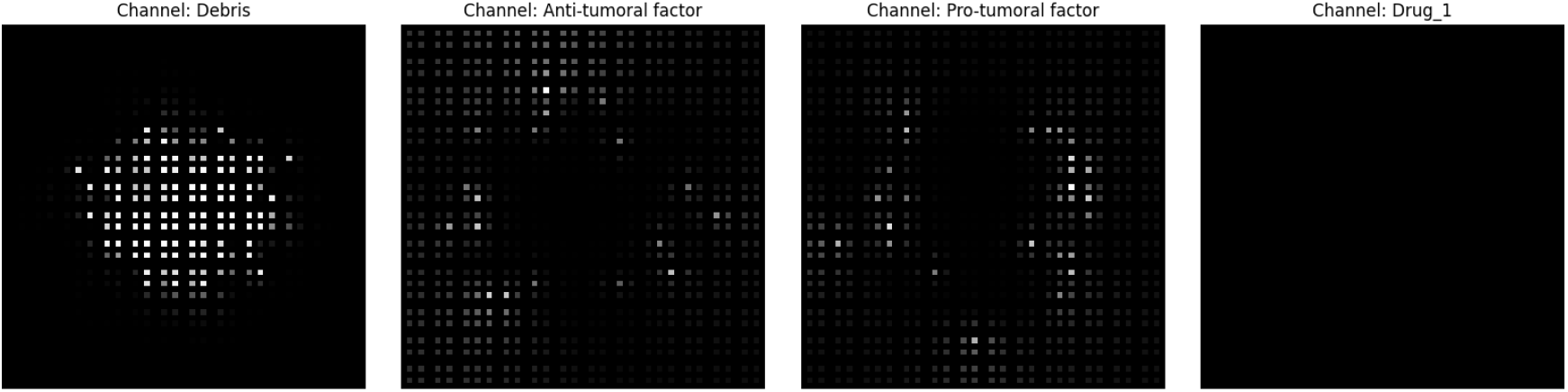
Representation of different channels; each channel represents a substrate.

An episode is truncated when the maximum limit of around 200 steps is reached, but it is also terminated when there are no more tumor cells in the environment.

To solve our control problem, namely to kill as many cancer cells as possible while reducing drug use, we chose Soft Actor-Critic (SAC) [7], which is a well-known RL algorithm designed for continuous control tasks.

We selected SAC for its sample efficiency, meaning its ability to learn effectively from a limited number of interactions with the environment. Additionally, as a deep RL method, SAC can process various types of input data, including images, by using Convolutional Neural Networks [14].

## 3 Results

We proposed a simplified TME simulation, and we used SAC to control it. We now describe our results when applying PhysiGym in this specific scenario. As seen in figure 4, our RL policy is able to learn how to dose the drug at each time step to eradicate cancer cells while minimizing the amount of drug used. We show the discounted cumulative return in figure 4.

**Figure 4.**
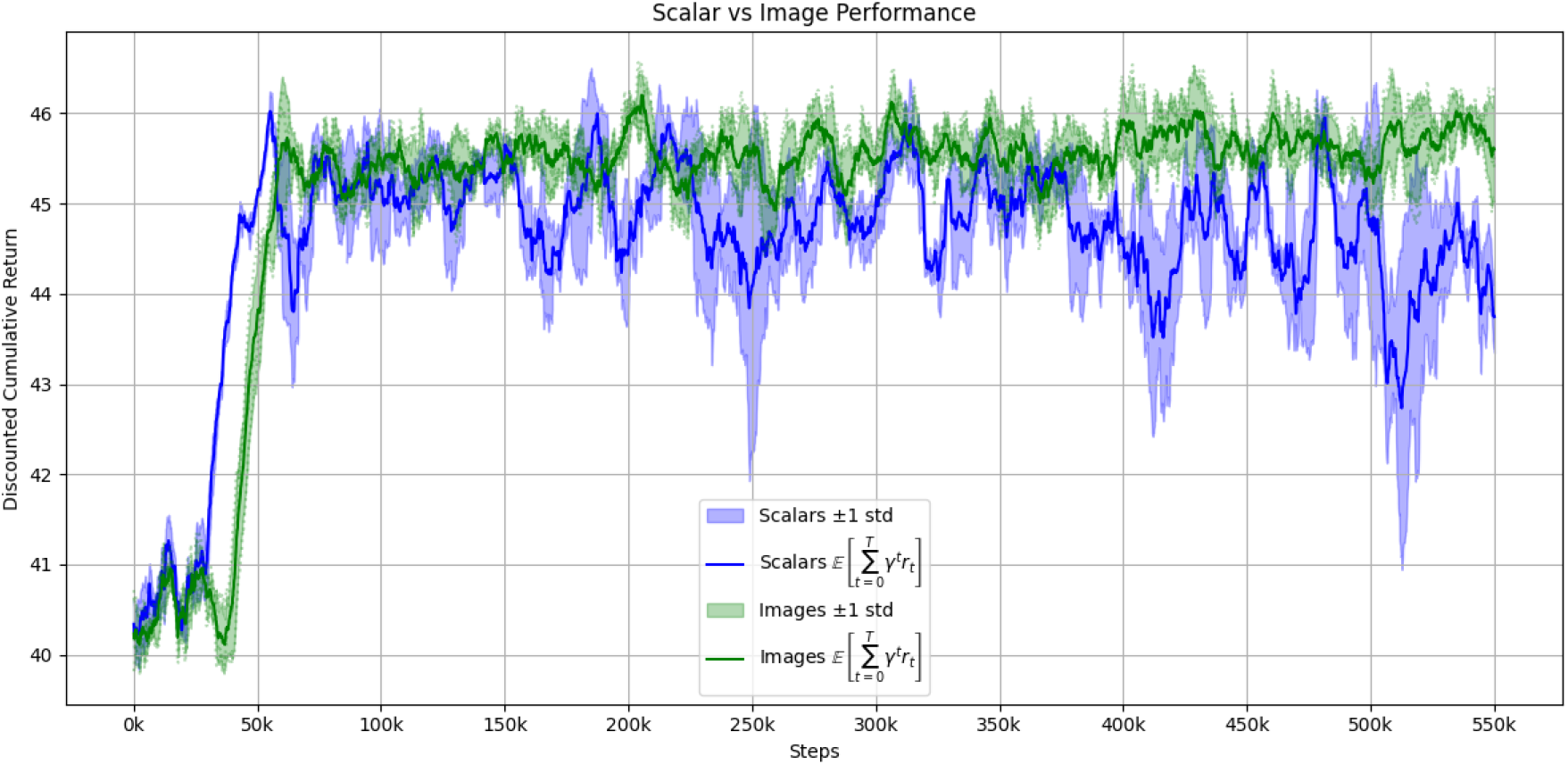
Discounted cumulative return (Y-axis) versus cumulative interaction steps with the environment (X-axis) across three different random seeds. The green line represents the expected discounted cumulative return for the image-based state space (*multi-channel cells & substrates*), while the blue line represents *scalars*. The lighter green (respectively blue) shading indicates *±*1 standard deviation around the mean for *multi-channel cells & substrates* (respectively *scalars*).

We compare two different state spaces: *multi-channel cells & substrates* and *scalars*.

Across different seeds, using the image-based state space (*multi-channel cells & substrates*) results in a more consistent performance, with lower variance compared to the scalar state space. Furthermore, this representation yields a higher expected discounted cumulative return overall. A divergence between the two state spaces is observed after 500k steps, where *multi-channel cells & substrates* significantly outperform *scalars*. The image-based state space exhibits a lower standard deviation, indicating a more robust and reliable representation compared to *scalars*.

For both state spaces, we observe different mean episodic lengths. Each state space leads to different treatment regimes.

Although the difference in discounted expected cumulative return after hundreds of thousands of steps is slight, as seen in figure 4, there is a significant difference in episodic length between the two state spaces (figure 5). We also see a greater standard deviation for the image-based state space compared to the scalars in terms of episodic length, while in terms of the discounted expected cumulative return, we observe the opposite. Taken together, these findings imply that using the *multi-channel cells & substrates* state space allows eliminating the tumor in fewer steps, adjusting the treatment more robustly compared to the strategy that only uses the *scalars* state space.

**Figure 5.**
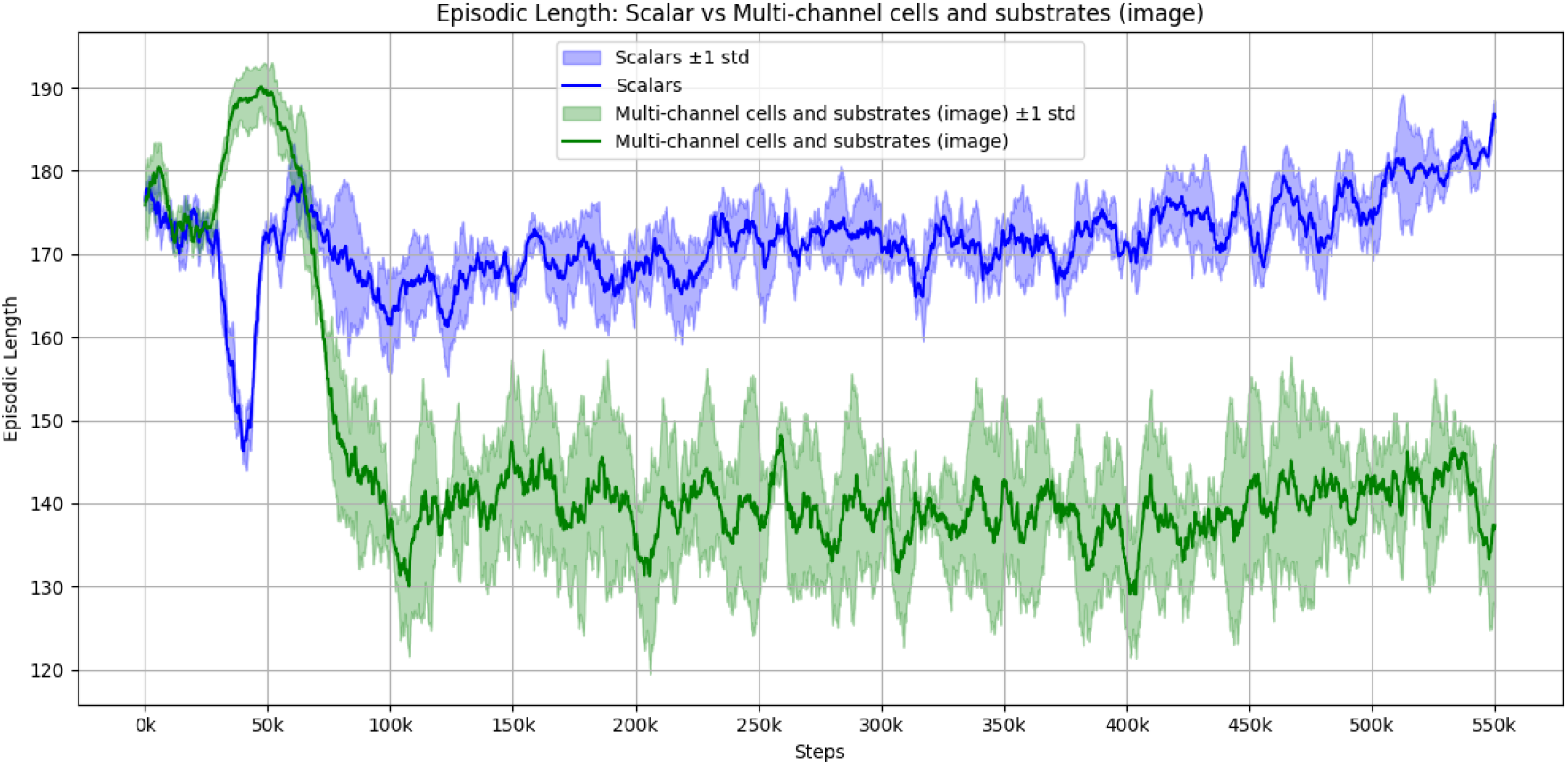
Episodic length (Y-axis) versus cumulative interaction steps with the environment (X-axis) across three different random seeds. The green line represents the mean episodic lengths for *multi-channel cells & substrates* state space, while the blue line represents *scalars*. The lighter green (respectively blue) shading indicates 1 standard deviation around the mean for *multi-channel cells & substrates* (respectively *scalars*).

For the two different state spaces, almost the same amount of drug was administered (cf. figure 6), but in terms of discounted cumulative return, the *multi-channel cells & substrates* state space performs better (cf. figure 4). For both state spaces, we can observe two distinct phases:

**Figure 6.**
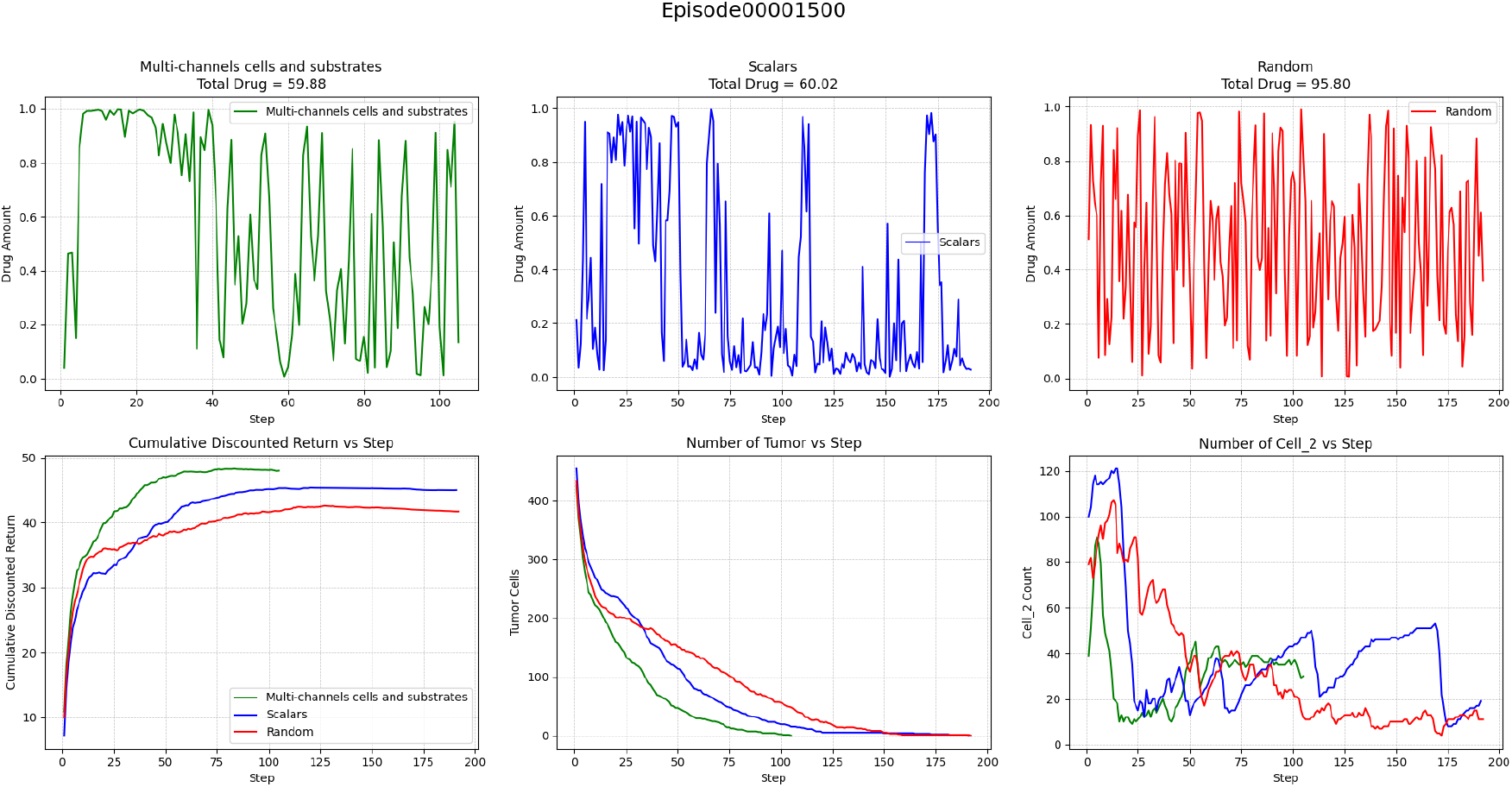
Representations of drug administration and outcomes for three dynamic treatment regimes at one specific episode: *multi-channel cells & substrates* (green, total drug = 59.88), *scalars* (blue, total drug = 60.02), and random policy (red, total drug = 95.80). Top row: drug amount per step for each regime. Bottom row: (left) cumulative sum of discounted cumulative return, (middle) number of tumor cells, and (right) number of `cell _2` cells over time.

- **Phase 1** refers to the initial steps, where `cell_1` changes into `cell_2`. The learned policy tries to administer as much drug as possible to convert cell 2 back into cell *±*1. This strategy appears effective because, in the early steps, there is greater pressure due to a higher number of cancer cells at the start.
- **Phase 2** refers to the remaining steps. This regime continues until the killing rate of `cell_1` exceeds the division rate of `tumor_cells`. Based on the environment dynamics, there are now enough `cell_1` to kill the remaining tumors even if some `cell _1` still convert into `cell_2`. This is explained by the fact that the total killing effect over time is much greater than:
  – the pro-inflammatory factor from `cell_2`,
  – the pressure causing `cell_1` to become `cell_2`,
  – the division rate of the remaining tumor cells.

Finally, the *multi-channel cells & substrates* state space provides more detailed insights, enabling a better treatment strategy that drastically reduces tumor size in fewer steps and gives a better discounted cumulative return, which is why the episodes are shorter.

## 4 Discussion

Our experiments demonstrate that PhysiGym enables RL policies to effectively control a simplified tumor microenvironment by integrating ABMs into a standardized framework. The results highlight that the choice of state representation shapes the learning process and the resulting treatment strategies. In particular, the image-based representation of cells and substrates (*multi-channel cells & substrates*) consistently outperformed the scalar representation, leading to higher cumulative returns and shorter episode lengths. This suggests that retaining spatial information allows the RL agent to exploit biologically meaningful features that would otherwise be lost in aggregated statistics. Such findings indicate that PhysiGym can serve as a platform to investigate how the system’s representation can impact therapeutic control strategies in complex biological environments.

While encouraging, these results are constrained by the simplicity of the underlying model and the chosen reward function. Real tumor-immune interactions involve greater heterogeneity, stochasticity, and multiscale processes, 3D-model, and may require moving from 2D to 3D models, which can further challenge the robustness of learned policies. Extending PhysiGym to richer biological models will therefore be essential to assess the generalization capacity of these models. Another limitation is that our experiments relied on a single RL algorithm (SAC). Although SAC is well-suited for continuous control, future work should investigate whether alternative approaches can lead to further improvements.

Beyond technical aspects, PhysiGym raises important research directions at the interface of biology and deep reinforcement learning. From a biological perspective, it provides a controlled testbed to hypothesize about potential treatment strategies, to be later tested in-vitro, ex-vivo or in-vivo in pre-clinical models and, ultimately, in clinical trials. From a computational perspective, PhysiGym enables systematic benchmarking of RL algorithms on structured, biological environments. These opportunities highlight PhysiGym’s potential to bridge simulation-based biology and decision-making AI in a reproducible and extensible manner.

## 5 Conclusion

PhysiGym provides a novel framework that bridges reinforcement learning with agent-based biological simulations, enabling the study of dynamic treatment regimes in silico. Our results show that leveraging spatial representations improves policy performance, underscoring the importance of biologically meaningful state spaces. By making PhysiGym open-source and extensible, we aim to foster collaboration between computational scientists and biologists, paving the way toward AI-driven strategies for personalized cancer therapy. We anticipate that PhysiGym will serve as a valuable testbed for both communities to advance decision-support systems that could translate into clinical practice in the future.

## 6 Code Management and Availability

PhysiGym is an open-source project, and contributions from the community are encouraged. New Physi-Cell models can be integrated by defining the required cell types and interactions, ensuring compatibility with PhysiGym’s step-based execution, exposing action and observation spaces in Python, and wrapping the model with a Gymnasium-compatible environment. Source code, examples, and documentation are available at https://github.com/Dante-Berth/PhysiGym.

Installation and troubleshooting how-tos, a modeling tutorial, and a RL tutorial, and a reference manual can be found at [https://github.com/Dante-Berth/PhysiGym/tree/main/man].

## 8 Acknowledgments

A. B., V.P., and E. R. acknowledge funding from Region Occitanie and INSERM, Projet Emergence REACT. E. B. received support from a Chateaubriand Fellowship of the Office for Science & Technology of the Embassy of France in the United States. This study has been partially supported through the grant EUR CARe N°ANR-18-EURE-0003 in the framework of the Programme des Investissements d’Avenir and an Eiffel Excellence doctoral fellowship to M. H. This research was also supported in part by Lilly Endowment, Inc., through its support for the Indiana University Pervasive Technology Institute.

